# AMP-activated protein kinase: A key enzyme to manage nutritional stress responses in parasites with complex life cycles

**DOI:** 10.1101/2020.04.08.032284

**Authors:** Tamara Sternlieb, Alejandra C. Schoijet, Patricio D. Genta, Guillermo D. Alonso

## Abstract

*Trypanosoma cruzi*, the etiological agent of Chagas disease, has a digenetic life cycle. In its passage from the insect vector to the mammalian host, and vice versa, it must be prepared to cope with abrupt changes in environmental conditions in order to survive. Sensing and signaling pathways that allow the parasite to adapt, have unique characteristics with respect to their hosts and other free-living organisms. Many of the canonical proteins involved in these transduction pathways have not yet been found in the genomes of these parasites, because they present divergences either at the functional, structural and/or protein sequence level. All of this makes these pathways promising targets for therapeutic drugs.

The AMP-activated protein kinase (AMPK) is a serine/threonine kinase activated by environmental stresses that results in reduction of ATP and increase of AMP levels. Thus, AMPK is regarded as a fuel gauge, functioning both as a nutrient and an energy sensor, to maintain energy homeostasis and, eventually, to protect cells from death by nutrient starvation.

In the present study, we report the characterization of AMPK complexes for the first time in *T. cruzi* and describe the function of TcAMPK as a novel regulator of nutritional stress in epimastigote forms. We demonstrate that this complex possesses specific AMPK kinase activity in epimastigotes, which is inhibited by Compound C and is modulated by carbon source availability. In addition, TcAMPKα2 subunit has an unprecedented functional substitution (Ser x Thr) at the activation loop and its overexpression in epimastigotes led to higher autophagic activity during prolonged nutritional stress. Moreover, the over-expression of the catalytic subunits resulted in antagonistic phenotypes associated with proliferation. Together, these results point to a role of TcAMPK in autophagy and nutrient sensing, key processes for the survival of trypanosomatids and for its life cycle progression.

## Introduction

Chagas disease, also known as American trypanosomiasis, is a potentially life-threatening illness endemic in Latin-American countries. This silent illness is conventionally caused when insects from the *Reduviidae* family transmit the protozoan parasite *Trypanosoma cruzi* to humans.

The World Health Organization (WHO), in the early 2000s, had 17 Neglected Tropical Diseases (NTDs) in its portfolio, including Chagas disease among them. Subsequently, in 2016 this list was expanded, adding three new groups of diseases to currently include 20 groups of NTDs (1). All NTDs are characterized by a high distribution in the world due to the increased number of infected people, even in non-endemic countries. Overall, NTDs were estimated to affect about 2 billion people at the turn of the millennium, with a collective burden of disability-adjusted life years that was equivalent to HIV/AIDS, tuberculosis, or malaria (2).

In addition to the widespread of these illnesses, very few drugs are annually developed for NTDs. Given that most NTDs affect low income groups and prices of the treatments could force these private entities to lower gains, drug developing companies show low interest in this line of products (3). For more than 50 years, Benznidazol and Nifurtimox have been used for Chagas disease treatment, both during its acute and chronic stages. However, their efficacy is limited and both drugs cause several side effects, leading to discontinuation of treatment.

Taking into account the above mentioned situation, the development of new technologies, higher government and industrial involvement and more scientists committed to basic investigation, are key to find an effective treatment and cure for the NTDs. In this sense, signal transduction pathways in trypanosomatids could be considered as an Achilles’ heel, since they are essential to recognize environmental fluctuations and allow these parasites to respond accurately through cellular changes. On this matter, advances in knowledge on the differences of these pathways between the parasite and mammalian cells might allow the identification of rational targets for the development of safe and more effective drugs for Chagas disease treatment (3).

Signal transduction plays a key role in regulating important functions in both multicellular and unicellular organisms and largely controls the manner in which cells respond to stimuli. *Trypanosoma cruzi* has a complex life cycle involving four main morphogenetic stages. Briefly, the life cycle of this parasite involves two intermediate hosts (triatomine insects and mammals, including man) and four well-defined morphological and functional developmental stages: epimastigotes, metacyclic trypomastigotes, amastigotes and bloodstream trypomastigotes. The epimastigote forms replicate in the midgut of the insect host and develop into non-replicative metacyclic trypomastigote forms. When the insects feed on blood, they release in their excreta metacyclic trypomastigotes that penetrate the body of the mammalian host through the wound and are able to invade cells. Within the host cells, the parasite differentiates into the replicative amastigote form. After multiplication, the amastigotes differentiate into bloodstream trypomastigotes that are released into the circulatory system, infecting new cells.

Various second messenger pathways are involved in the regulation of the cell proliferation and the passage from epimastigotes to metacyclic trypomastigotes, a process known as metacyclogenesis. Although several factors have already been identified, the mechanisms and events involved within the onset of metacyclogenesis remain to be elucidated.

In addition, these parasites are also of biological interest since they possess structures and organelles that are not found in mammalian cells. Therefore, understanding of the unique pathways in this pathogen may lead to the development of novel therapeutic agents. In particular, targeting metabolic pathways in the parasite for rational drug design represents a promising research field.

Cells with a sufficient energy supply maintain a balance between the adenosine phosphate species of 10:1 for ATP:ADP and 100:1 for ATP:AMP, thanks to the interconversion between them, executed by ATPases, ATP synthases and Adenylate Kinases (4). From the nature of these enzymatic reactions, when the energy status is affected and ATP diminishes, the AMP concentration increases as the square of the ADP concentration. This explains why most energy sensing mechanisms in the cell depend on AMP detection and activation. One of these mechanisms is the AMP-activated protein kinase (AMPK) complex.

The cellular energy homeostasis can be affected by different kinds of stresses, such as nutritional, oxidative, heat shock, amongst others. Enzymes like the AMPK can respond in a very sensitive manner to the imbalances caused by these stresses and thus shift cellular anabolic activity to a more catalytic one, which replenishes the ATP levels.

The AMPK complex exists as a heterotrimer, composed of an alpha catalytic subunit and two regulatory subunits, beta and gamma. The α subunit contains a kinase domain as well as a regulatory domain that inhibits the enzyme in the absence of AMP (5). The β subunit acts as a scaffold for the other components while also modulating the localization of the complex (6–8), and the γ subunit is involved in AMP binding (9). With very few exceptions, every eukaryotic organism expresses at least one AMPK complex. Amino acid sequence of the subunits vary between different organisms but conserve a domain architecture that allows the execution of ancient functions. Yet, small mutations in some codons also led to neofunctionalization of the orthologs and paralogs, diversifying the role of this complex between organisms and tissues. The AMPK yeast ortholog, SNF1, is known for its role in the inhibition of glucose gene repression. In higher eukaryotes, AMPK phosphorylates enzymes of the isoprenoid synthesis and fatty acid synthesis pathways to inhibit these metabolisms and switch the cell to the use of stored or alternative carbon sources, such as fatty acids (10). AMPK can also phosphorylate transcription factors, inducing the expression of proteins involved in several stress response pathways. Through all these functions, this complex has been established as an essential regulatory hub for the cell.

Parasitic organisms fully depend on the environmental signals in their hosts to progress in their life cycle. The rapid passage through these environments also exerts several kinds of stresses that parasites must overcome to survive. Molecular stress sensors, such as AMPK, can play key roles in these processes. Recent studies have reported that *T. brucei* expresses an AMPK complex (TbAMPK) with two possible catalytic α subunits. TbAMPK is involved in the expression of membrane proteins in response to nutritional stress and in the transition from the slender bloodstream form to the quiescent stumpy (11,12). Also, Saldivia et. al. demonstrated that this complex is responsible for the growth arrest observed when parasites are treated with AMP analogs. TbAMPK complex containing the α1 subunit is also capable of responding to AMP:ATP imbalances produced by mitochondrial depolarization, and once activated it phosphorylates proteins in the glycosomes, the organelles nucleating the first steps of glycolysis in these organisms (13). However, some of the conventional roles of the AMPK complexes may not be conserved in the trypanosomes. Autophagy, a process that allows cells to consume their own components to provide nutrients and overcome stress, and which involves proteins activated by AMPK phosphorylation in many organisms, does not appear to depend on AMPK activation in *T. brucei* (14). All these recent discoveries point to novel roles of AMPK in trypanosomatids, associated with their complex life cycle and ability to differentiate in response to environmental cues.

In this work, we identify the corresponding genes for the TcAMPK subunits, confirm the activity of the TcAMPK complex in the epimastigote stage and demonstrate its modulation under nutritional stress. Finally, we show that the TcAMPK complex containing the α2 subunit is involved in autophagy when parasites are deprived of a carbon source, being this the first report where an AMPK complex from trypanosomatids shows to be involved in autophagy.

## Methods

### Parasite cultures

*T. cruzi* epimastigotes of the CL Brener strain were cultured at 28 °C for 7 days in liver infusion tryptose (LIT) medium (5 g/l liver infusion, 5 g/l bacto-tryptose, 68 mM NaCl, 5.3 mM KCl, 22 mM Na2HPO4, 0.2% (w/v) glucose, and 0.002% (w/v) hemin) supplemented with 10% (v/v) FCS, 100 U/ml penicillin and 100 mg/l streptomycin. Cell viability was assessed by direct microscopic examination. When assessing growth curves, duplication times were calculated as follows:

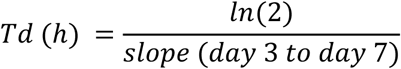

Where the slope between day 3 and 7 was obtained by a linear regression on the quantification of parasites in a Neubauer chamber over the number of days.

### Generation of TcAMPK-subunits overexpressing cell lines in *T. cruzi epimastigotes*

The full-length genes of every TcAMPK subunit were amplified using the following primers: for TcAMPKα1 TcAMPKα1-Fw-HindIII 5’- GGATCCATGAGTCAGAAGTTTGGCCCCTA-3’ and TcAMPKα1-Rv-pRIBOHA-XhoI 5’- CTCGAGTTAAGCGTAATCTGGAACATCGTATGGGTATTCGTCTGGTCCAAGAGAGGAA-3’. For TcAMPKα2 TcAMPKα2-Fw-HindIII 5’- AAGCTTATGCATTCCAGGCGGGATGTT-3’ and TcAMPKα2-Rv-pRIBOHA-XhoI 5’- CTCGAGTTAAGCGTAATCTGGAACATCGTATGGGTAACCCATTCGATGAACGAGC GTC-3’. For TcAMPKβ TcAMPKβ-Fw-HindIII 5’- GGTACCATGGGCCAACAAAATGCCAGGGA-3’ and TcAMPKβ-Rv-pRIBOHA-XhoI 5’- CTCGAGTTAAGCGTAATCTGGAACATCGTATGGGTACCCGTTCGGAGCTCCCATTCT-3’. For TcAMPKγ TcAMPKγ-Fw-HindIII 5’- AAGCTTATGCGTCGCACGAGTGCCTTTGC-3’ and TcAMPKγ-Rv-pRIBOHA-XhoI 5’- CTCGAGTTAAGCGTAATCTGGAACATCGTATGGGTATTTTTGTGCGTTGCCGTCATT -3’. The PCR products, now containing an hemagglutinin tag at their C-terminal end, were then cloned into pGEM-T Easy plasmid, the sequence identity was confirmed by DNA sequencing, and subcloned into the pRIBOTEX plasmid (15). *T. cruzi* epimastigotes of CL Brener strain were transfected with the pRIBOTEX constructs as described previously (16). Stable cell lines were achieved after 60 days of treatment with 500 µg/ml G418 (Gibco BRL, Carlsbad, CA) and the transgenic condition was confirmed by western blot analyses.

### Yeast transformation and functional complementation

Conditional *Saccharomyces cerevisiae* yeast mutant strains were generously provided by Dr. Martin C. Schmidt, from the University of Pittsburgh (17,18). The same pGEM-T Easy constructs with each TcAMPK subunit were used to subclone into the p416 yeast expression plasmid. Yeasts were transformed following the “Quick and Dirty” protocol (19). Briefly, a fresh yeast inoculum was mixed with 100 μl of transformation mix (200 μl of 2 M sterilized LiAc, 800 μl of sterilized 50% PEG-3350, 7.7 μl of 14 M 2-Mercaptoethanol) freshly prepared, 3 μl of denaturalized salmon sperm DNA (10 mg/ml) and 1 μg of plasmid DNA or distilled water as control. The mix was incubated at 37 °C for 30 min while mixing. After centrifugation at 3000 rpm for 5 min, the pellet was recovered and resuspended in 100 μl of sterile water. The transformed yeasts were plated on selective media (lacking uracil) and were grown for 16 to 72 h at 30 °C. Individual colonies were tested for heterologous protein expression. A positive clone was grown overnight at 30 °C in YPAD media (yeast extract 10 g/l, peptone 20 g/l, glucose 20 g/l, adenine sulfate 0.04 g/l) up to an OD_600_ of 2. Serial dilutions of the culture were plated on selective media without uracil (0.17% (w/v) yeast nitrogen base (without amino acids and ammonium sulphate) and 0.5% (w/v) ammonium sulphate, supplemented with the corresponding amino acid mixture) and containing either glucose (2% w/v) or raffinose (2% w/v) as the carbon source. Plates were incubated at 30 °C between 3 to 5 days after which growth was photographed.

### Protein kinase assay

*In vitro* kinase reactions were developed in a final volume of 50 μl containing 0.02 mM [γ-^32^P]ATP (1 µCi per tube, Perkin Elmer, Massachusetts, USA), 50 mM Tris-HCl, pH 7.0, 0.1 mM NaCl, 0.1 mM EDTA, 0.5 mM dithiothreitol, 5 mM MgCl_2_ and 100 μM SAMS peptide (HMRSAMSGLHLVKRR, ab120182, Abcam) as AMPK substrate. The reaction was initiated by adding 50 μg of epimastigote protein extract and was incubated with shaking at 30 °C for 10 min. As enzyme blank control, tubes where protein extract was replaced by lysis buffer were added. The reaction was stopped by immediate immersion in an ice bath and spotting on P81 Whatman filter paper. Unreacted ATP was removed washing the filter papers 3 times with 1% phosphoric acid for 7 min. After the final wash, the filters were quickly dried with ethanol, placed in polistor tubes with 2 ml of Ultima gold™XR liquid scintillation cocktail (Perkin Elmer, Massachusetts, USA) and counted in a scintillation counter. AMPK activity was calculated as phosphorus incorporation subtracting the average counts per minute (cpm) of the enzyme blank controls from the average cpm of protein extract samples. For Dorsomorphin (CC) (#P5499, Sigma Aldrich, St. Louis, Missouri, United States or ab120843, Abcam, Cambridge, United Kingdom) treatment protein extracts were incubated in an ice bath with the 1 µM final concentration of the inhibitor for 10 min previous to the addition of the mix containing the SAMS.

### Cell extracts and western blotting

To obtain *T. cruzi* extracts, 10^8^ epimastigotes were harvested by centrifugation at 1500 g for 10 min and washed two times with phosphate-buffered saline (PBS). Cell pellets were then resuspended in a lysis buffer (50 mM Tris-HCl buffer, pH 7.5; 14 mM 2-Mercaptoethanol, PMSF and E64 as proteases inhibitors) and lysed by six cycles of freezing in liquid N_2_ and thawing at 4 °C.

For western blotting analysis, proteins were solved in 10% (w/v) SDS-polyacrylamide gel electrophoresis as described by Laemmli (1970) and electrotransferred to Hybond-C membranes (Amersham Pharmacia Biotech, Piscataway, USA). The membranes were blocked with 5% (w/v) non-fat milk or 5% (w/v) BSA suspension in 0.05% TBS-Tween for at least 3 h. Blocked membranes were then incubated overnight with a 1:1000 dilution of Phospho-AMPKα (Thr172) (40H9) Rabbit mAb (#2535, Cell Signaling, Massachusetts, USA). Detection was carried out by incubating with a 1:5000 dilution of a goat anti-rabbit antibody conjugated to peroxidase (Sigma Aldrich). For tagged proteins detection, membranes were incubated for at least 1 h with a 1:1000 dilution of a high affinity anti-HA antibody from rat IgG1 (#11867423001, Roche Applied Science, Penzberg, Germany). Detection was carried out by incubating with a 1:4000 dilution of a rabbit anti-rat antibody conjugated to peroxidase (A5795, Sigma-Aldrich). The membranes were then developed with the ECL Plus™ Western blotting detection system (PerkinElmer Life Sciences, Massachusetts, USA).

For AICAR (ab120358, Abcam, Cambridge, United Kingdom) treatment, epimastigotes were incubated for 30 min with a final concentration of 1 mM AICAR in LIT media. For CC and AICAR simultaneous treatment, epimastigotes were incubated with both compounds (10 µM of CC) for 1 h.

When specified, aliquots of the same protein extract were separated and treated with 200 U of Lambda Phage Phosphatase (#P0753S, New England Biolabs, Ipswich, Massachusetts, United States), as specified by the manufacturer, for different time periods. The reaction was stopped by boiling in Laemmli buffer.

### Proteomic analysis

After 10% SDS-PAGE of epimastigotes protein extracts, the section of the gel between 70 kDa and 100 kDa was manually cut and preserved in an eppendorf tube at -80 °C until delivery for mass spectrometry analysis. Mass spectrometry analysis was carried out at Centro de Estudios Químicos y Biológicos por Espectrometría de Masa (CEQUIBIEM), Argentina, in a Q Exactive HESI-Orbitrap coupled to a nano HPLC Easy-nLC 1000 (Thermo Scientific). Resulting reads were matched to the *T. cruzi* proteome available at the UNIPROT proteome database.

### Indirect Immunofluorescence

For immunofluorescence, cells were fixed with 4% paraformaldehyde in PBS for 20 min. Next, the cells were washed twice in Dulbecco’s PBS, pH 7.2, adhered to poly-L-lysine-coated coverslips, and permeabilized for 10 min with 0.3% Triton X-100. Cells were incubated for 30 min with 25 mM ammonium chloride and washed again with PBS, after which they were blocked for 20 min in 3% bovine serum albumin in PBS, pH 8.0, and incubated for 1 h with rat anti-HA high affinity monoclonal antibodies (Roche Applied Science, Penzberg, Germany) at 1:500. Cells were then washed in 0.05% TBS-Tween buffer, incubated with the secondary antibody, anti-rat Alexa 546 conjugate at 1:500, and mounted with Vectashield (Vector Laboratories, California, USA) containing 5 mg/ml DAPI. Cells were observed in an Olympus BX41 fluorescence microscope and images were captured.

### Autophagy monitoring by Monodansylcadaverine incorporation

Epimastigotes in exponential phase (2×10^7^ parasites per ml) were washed with PBS twice and resuspended in fresh LIT medium as control or PBS for starvation. Cultures were incubated at 28 °C for 17 h. Staining of autophagosomes with Monodansylcadaverine (MDC, #D4008, Sigma-Aldrich, St. Louis, Missouri, United States) was applied as in Munafó and Colombo (2001) (20). Briefly, after 16 h starvation of epimastigotes, MDC was added at 0.05 mM final concentration and incubation proceeded for 1 h at 28 °C. Epimastigotes were then washed two times with PBS and lysed in a buffer containing 10 mM Tris HCl pH 8.0 and 1% v/v Triton-X100. Lysates were distributed in triplicates in a 96 well plate and Ethidium bromide was added to the lysates at 0.2 μM per well. Fluorescence emission was measured at 380 nm excitation and 525 nm emission for MDC and 530 nm excitation and 590 nm emission for Ethidium bromide. Readings were quantified by fluorescence photometry in a Synergy HTX Multi-mode plate reader and normalization of the measures was calculated as MDC_525nm_/BrEt_590nm_ and expressed in arbitrary units.

## Results

### *In silico* identification of TcAMPK subunits and evolutionary analysis

Previously, Clemmens et. al. identified the beta and gamma AMPK subunits in *T. brucei*, and Salvidia et. al. identified both isoforms of the alpha subunits in the same organism ^11,12^. We used those protein sequences as baits for a BLAST search at the Tritrypdb database (https://tritrypdb.org/tritrypdb/) to find the corresponding orthologs in *T. cruzi*. The protein sequences retrieved from the CL Brener Esmeraldo like strain for the beta and gamma subunits (IDs TcCLB.504427.50 for the TcAMPKβ and TcCLB.503841.20 for the TcAMPKγ) have 47.78% and 51.92% sequence identity with *T. brucei*, respectively. Whereas the alpha subunit orthologous genes in *T. cruzi* (TcCLB.506679.80 for TcAMPKα1 and TcCLB.510329.210 for TcAMPKα2) have around 58 % sequence identity with their respective *T. brucei* orthologs (Figure 1B). Interestingly, the paralogs only share around 30% identity between them.

**Figure 1.**
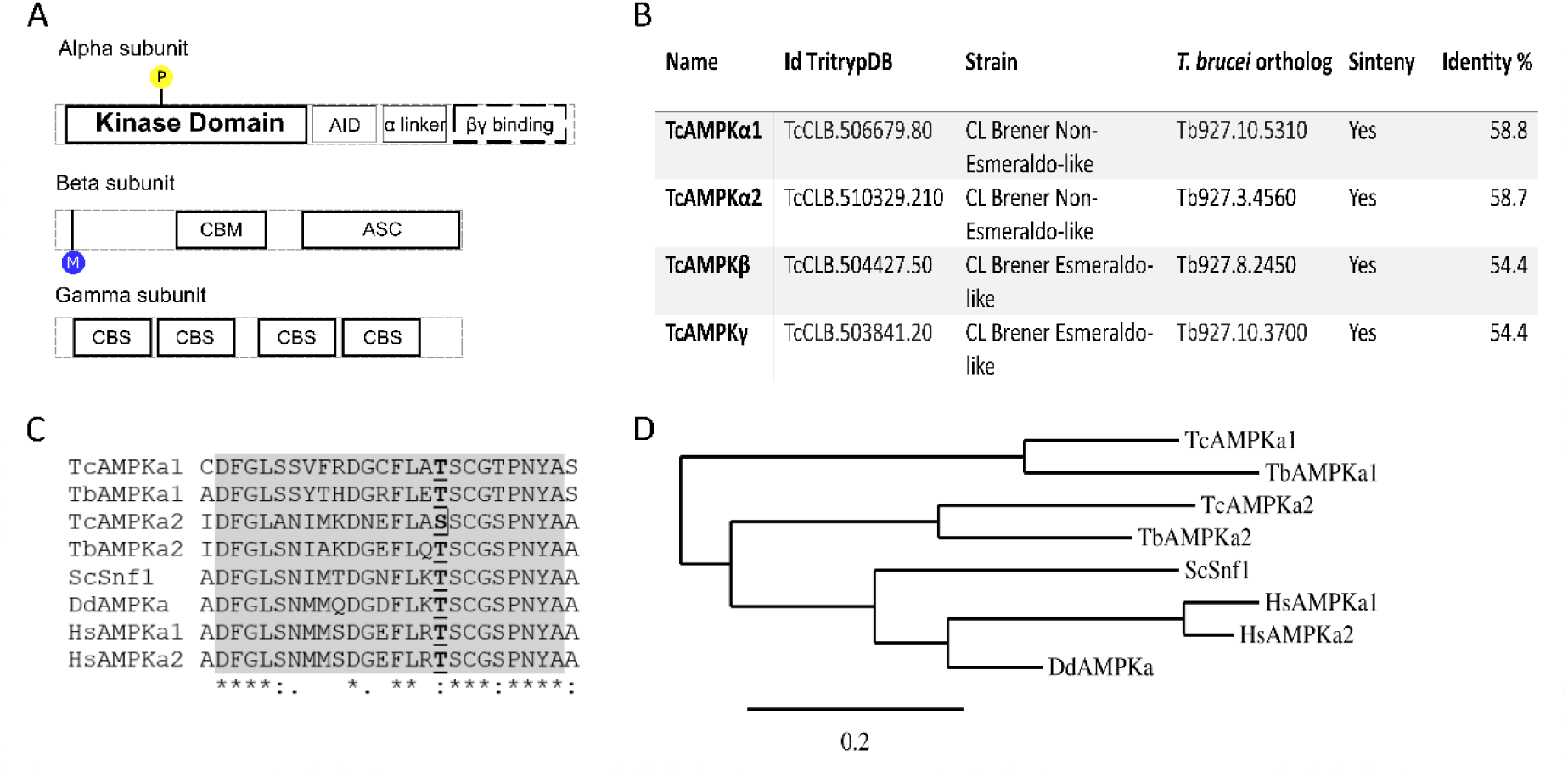
AMPK is partially conserved in *T. cruzi*. (A) Protein domains of every AMPK subunit. Domains with a dark border line are conserved in *T. cruzi*, while domains with dashed lines borders are only partially conserved. AID and α linker domain could not be identified in *T. cruzi* catalytic subunits. (B) Table of IDs for TcAMPK subunits and the percentage identity with respect to their orthologous in *T. brucei*. (C) Section of the multiple alignment of the alpha subunits of several eukaryotic organisms. The activation loop conserved region is painted in grey, and the Threonine (or Serine) residue phosphorylated during AMPK activation is in bold font and underlined. (D) Phylogenetic tree of the full-length multiple sequence alignment in C.

The regulatory subunits candidates show the expected protein domains to function in an AMPK complex (Figure 1A). Key amino acids for the subunits interactions remain conserved, and the protein domains are readily recognized by the NCBI Conserved Domain Search. However, the kinase catalytic domain is the only conserved region predicted in the alpha subunit candidates. The C-terminal portion of AMPKα usually presents an autoinhibitory sequence or domain (AIS or AID), a linker and a βγ-interaction domain. These C-terminal domains could not be recognized by any informatic tool on the TcAMPKα orthologs, although PROSITE could identify a sequence called “Kinase Associated domain”, of unspecified function. Another relevant trait of the alpha subunit in this kinase complex is the activation loop, which contains a conserved Threonine residue commonly known as Thr172, for its location in the *Rattus sp*. AMPKα protein sequence. This Thr must be phosphorylated to reach maximum levels of kinase activity. As an outstanding feature, TcAMPKα2 presents a Serine residue replacing the Threonine present at the activation loop, further emphasizing the differences between TcAMPKα1 and TcAMPKα2 subunits (Figure 1C). This phosphorylatable residue is highly conserved and TcAMPKα2 is the only coding sequence, to our knowledge, that presents this divergence.

Interestingly, in both *T. Cruzi* and *T. brucei*, AMPKα subunits share a low sequence identity between its paralogous genes, which is surpassed by the identity between orthologous sequences. This could mean that the gene duplication event occurred before the speciation of these two trypanosomatids, illustrated in the phylogenetic tree in Figure 1D. Future studies on the evolution of these genes could uncover neofunctionalization of these AMPK complexes in trypanosomatids.

### Functional complementation in yeast

To evaluate the functional capability of each of the putative TcAMPK subunits, we performed complementation assays in *S. cerevisiae* conditional mutant strains, which are alternatively deficient for alpha subunit (MSY1217), the three beta subunits (MSY557) or gamma subunit (MSY846). These mutants are unable to grow in media containing any carbon source other than glucose. The putative TcAMPK subunits were subcloned into p416 yeast expression vector as fusion proteins to a C-terminal HA-tag and used to transform the corresponding yeast strain. All TcAMPK subunits, TcAMPKα1-HA, TcAMPKα2-HA, TcAMPKβ-HA and TcAMPKγ-HA, were able to restore the capability of their specific conditional mutant to use raffinose as a carbon source (Figure 2). On the other hand, the same strains transformed with the empty vector only grew when glucose was added to the culture media, but not when it was replaced by raffinose. These results not only indicate that *T. cruzi* AMPK subunits are functional in yeast but also allow us to propose that *T. cruzi* AMPK conserves the main subunit functions present in *S. cerevisiae*. Western blots of protein extracts from transformed yeasts, revealed with anti-HA antibody, were performed to confirm the expression of each constructs (Figure 2, right panels).

**Figure 2.**
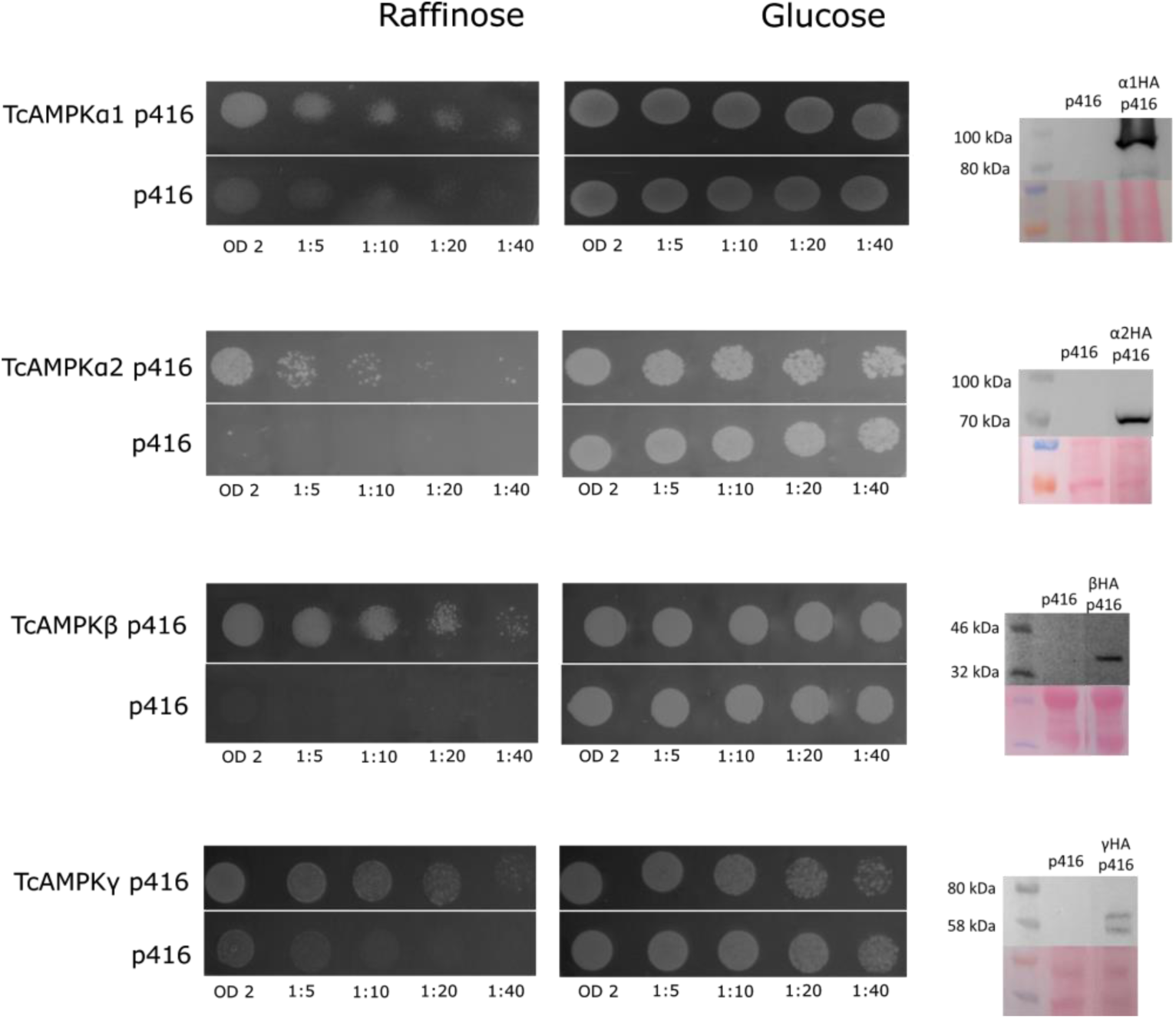
TcAMPK subunits can complement *S. cerevisiae* conditional mutants deficient in SNF1 subunits. Transformed yeast were plated in serial dilutions on minimal medium containing an Uracil drop-out and either glucose or raffinose as the only carbon source. Transformation of the conditional mutant yeasts with TcAMPK subunits restored the wild type phenotype, enabling the consumption of raffinose as a carbon source. The yeast containing the empty p416 plasmid could not grow on medium containing raffinose. Western blots at the right margin show the expression of the heterologous construct.

### Evaluation of TcAMPK activity in *T. cruzi* epimastigotes

Once we determined that TcAMPK subunits can act as functional components of an AMPK complex, capable of restoring the use of raffinose as a carbon source in conditional mutant yeasts, we decided to study the *in vivo* modulation of TcAMPK catalytic activity and initiate its biochemical characterization. To begin with, we established the assay conditions to measure AMPK specific kinase activity in epimastigote protein extracts by phospho transference of ^32^P from [γ-^32^P]ATP to the AMPK-specific substrate, SAMS (21,22). Figure 3A shows that, under the determined assay conditions, without the addition of SAMS in the reaction mix, only a basal kinase activity was observed. On the other hand, when SAMS was added as substrate, kinase activity increased significantly. As a further evaluation of the presence of a specific AMPK activity, we tested the effect of Dorsomorphin (also known as Compound C or CC, a potent, selective, reversible, and ATP-competitive inhibitor of AMPK) (23) on the SAMS phosphorylation capability of epimastigotes extract. Figure 3B shows a potent inhibitory effect of CC on the kinase activity, decreasing to basal levels similar to the activity observed in the absence of SAMS.

**Figure 3.**
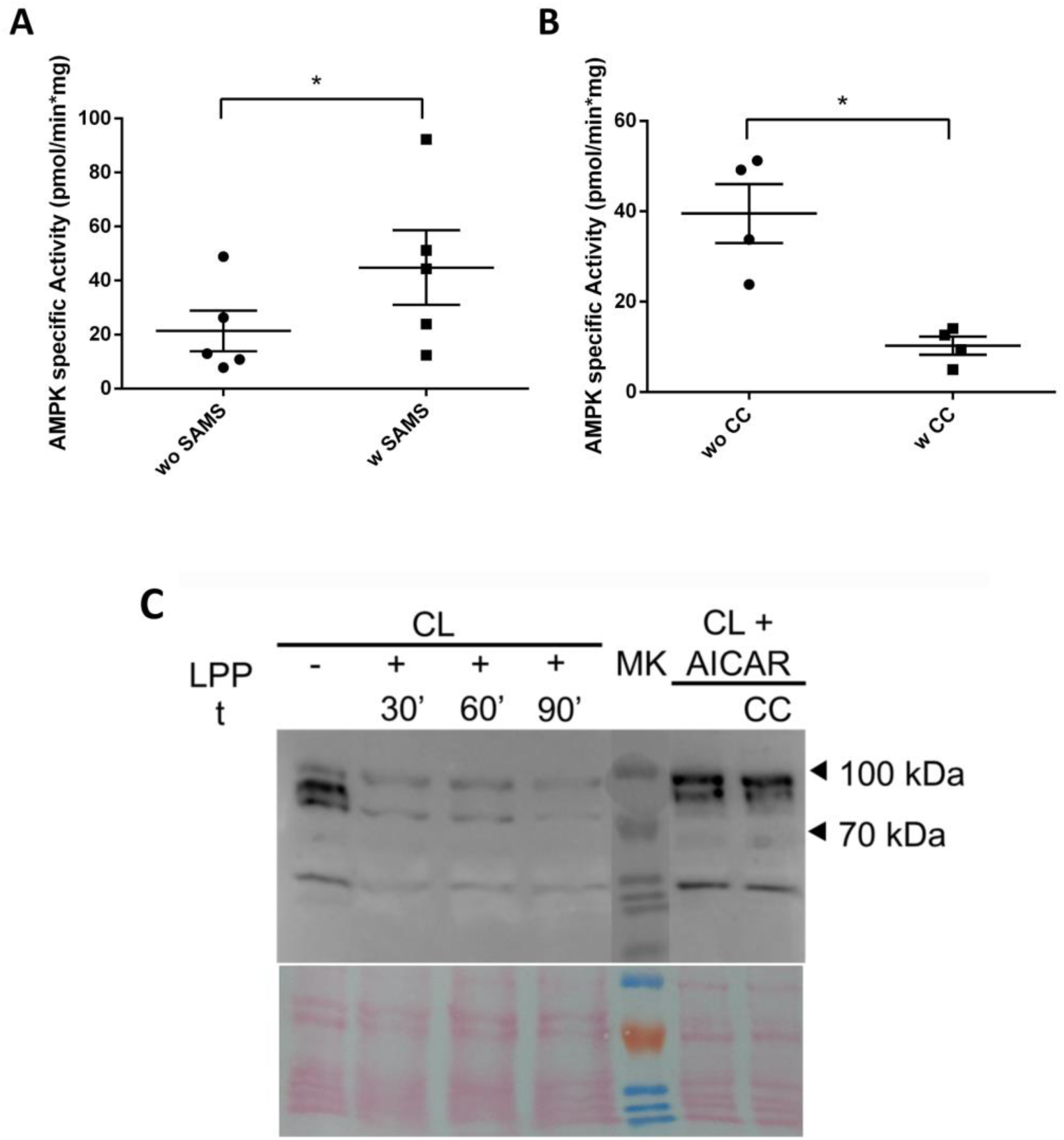
Kinase activity assays and detection of AMPKα subunit phosphorylation confirm *T. cruzi* epimastigotes express a functional AMPK. (A) Epimastigote whole protein extracts were tested for specific AMPK substrate, SAMS, phosphorylation by incorporation of 32P. Results show a significant increase in activity when SAMS is added to the mix (n = 5). (B) Addition of Compound C, an AMPK specific inhibitor, to the activity mix decreases activity to basal levels (n = 4). (C) Phosphorylation of the activation loop of AMPKα subunits was tested with Phospho-AMPKα (Thr172) antibody. Two bands of the expected weights for both TcAMPKα subunits were revealed. Lambda Phage Phosphatase treatment eliminated these markings. AICAR treatment, a specific AMPK activator, of epimastigotes cultures modified the intensities of the two bands. This effect was not altered by CC simultaneous treatment. All lanes contain equal mass of protein extract. Bottom panel shows the Ponceau staining as loading control. Representative assay from at least two experiments. CL: CL Brener epimastigote protein extract; LPP: Lambda Phage Phosphatase; MK: molecular weight marker; CC: Compound C; t: time. Error bars represent Standard Error of the Mean. Statistical test corresponds to paired t-test. * p value < 0.05.

Another tool that allows the visualization of the activation status of AMPK catalytic subunits is a commercial antibody developed against the phosphorylated state of the Thr172 in the activation loop. We tested this antibody against epimastigote protein extracts and obtained two reaction bands of the theoretical sizes predicted for the alpha subunits of *T. cruzi* (Figure 3C). To further confirm their phosphorylated status, we treated the protein extracts with Lambda Phage serine/threonine Phosphatase (LPP) for increasing time periods (30, 60 and 90 min). Throughout these incubation times, it was possible to observe that the bands’ intensity decreased. We also studied the AMPK phosphorylation state in response to specific treatments with AMPK activity modulators. We treated epimastigotes with AICAR, an AMP analog which is known to act as a specific AMPK activator (24), and observed a shift in the intensities of the phosphorylated bands. Incubation with both AICAR and CC didn’t change the previously observed shift. These results can be explained by the mechanisms of action of the two compounds. AICAR is an analog of AMP and, as such, it can activate AMPK by binding with the gamma subunit, inducing a conformational change that protects the Thr172 from dephosphorylation, while CC is a competitive inhibitor binding to the ATP site in the catalytic subunit. Hence, CC inhibits the kinase activity but doesn’t necessarily affect the phosphorylation status.

Gel regions surrounding the molecular weight between 70kDa - 100kDa of denaturing SDS-PAGE from protein extracts were excised and prepared for mass spectrometry analysis. Through this analysis, we have been able to detect the TcAMPKα2 and the phosphorylation in the Serine replacing the more canonical Threonine in the activation loop (Supplementary Figure 1). This result confirms the possibility of a catalytic subunit activation with a natural Ser for Thr substitution in an AMPK catalytic subunit.

### Catalytic activation of TcAMPK under nutritional stress

In many organisms, AMPK is a metabolic regulator that is activated when energy metabolites and nutrients are limited (25). Following this rationale, we hypothesized that TcAMPK could be a key enzyme sensing the different metabolic states that epimastigotes undergo during its passage through the insect gut. To further evaluate this hypothesis, *T. cruzi* epimastigotes were exposed to nutritional stress by incubation in Phosphate Buffered Saline (PBS) during seventeen hours at 28 °C. This condition has previously been used to study other processes related to nutritional stress in these parasites, such as autophagy (26,27). The AMPK catalytic activity after this treatment shows a mean increase of 2 times over the activity in the control condition. On the other hand, incubation in the same buffer supplemented with glucose 2% (w/v) prevents the enzyme activity increase. Protein kinase activity values were normalized to that obtained from parasites grown in LIT medium (Figure 4). These results demonstrate that TcAMPK can also be activated by the absence of a carbon source and suggest a possible role initiating metabolic responses to face prolonged nutritional stress in *T. cruzi* epimastigotes.

**Figure 4.**
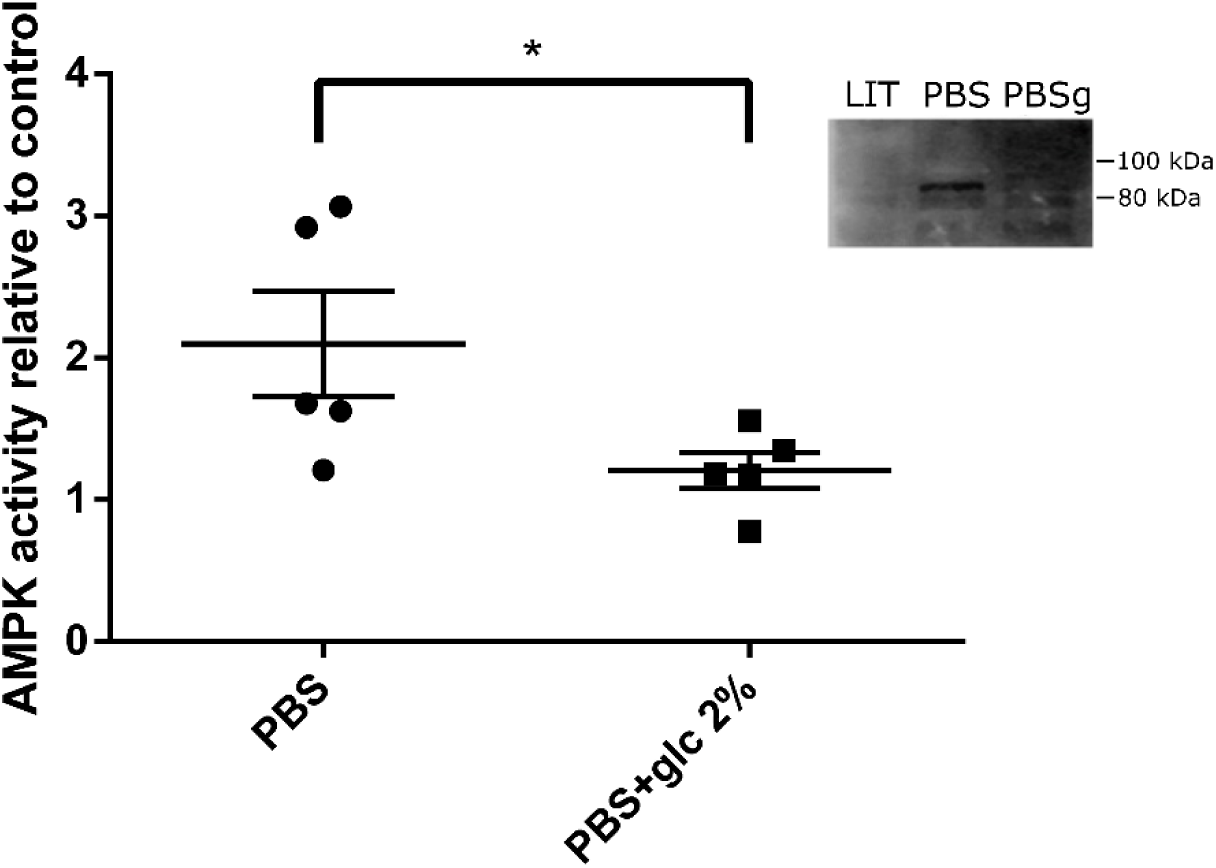
Nutritional stress increases AMPK activity in epimastigotes, and addition of glucose keeps AMPK activity at basal levels. Protein kinase activity was tested in the presence of SAMS and epimastigote whole protein extracts. Epimastigotes were incubated for 17 hs alternatively in LIT medium as control, PBS or PBS containing 2% (w/v) glucose. Measurements were normalized to LIT cultures and a paired t-test was performed to compare AMPK activity differences (n = 5). The inset image shows a western blot of epimastigote protein extracts under these same treatments revealed with the anti-Phospho AMPKα antibody. *p value < 0.05

### Effect of TcAMPKα isoforms overexpression on epimastigotes proliferation

To further reveal metabolic pathways that involve TcAMPK, both TcAMPKα1-HA and TcAMPKα2-HA constructs were subcloned into the pRIBOTEX expression vector and *T. cruzi* epimastigotes were then independently transfected with these plasmids. After the selection of stable transgenic lines, the expression and intracellular localization of these two proteins were investigated by western blot and indirect immunofluorescence using an anti-HA antibody (Figure 5 A and B). Western blots showed both overexpressed proteins of molecular weights between 70 kDa and 100 kDa, with TcAMPKα1-HA slightly heavier than TcAMPKα2-HA (Figure 5A, right margin). Interestingly, TcAMPKα1-HA appeared as a single band, while TcAMPKα2-HA appeared as two bands of very similar weight. This double band could suggest a post-translational modification affecting the protein migration. Further assays will be necessary to reveal the nature of this modification.

**Figure 5.**
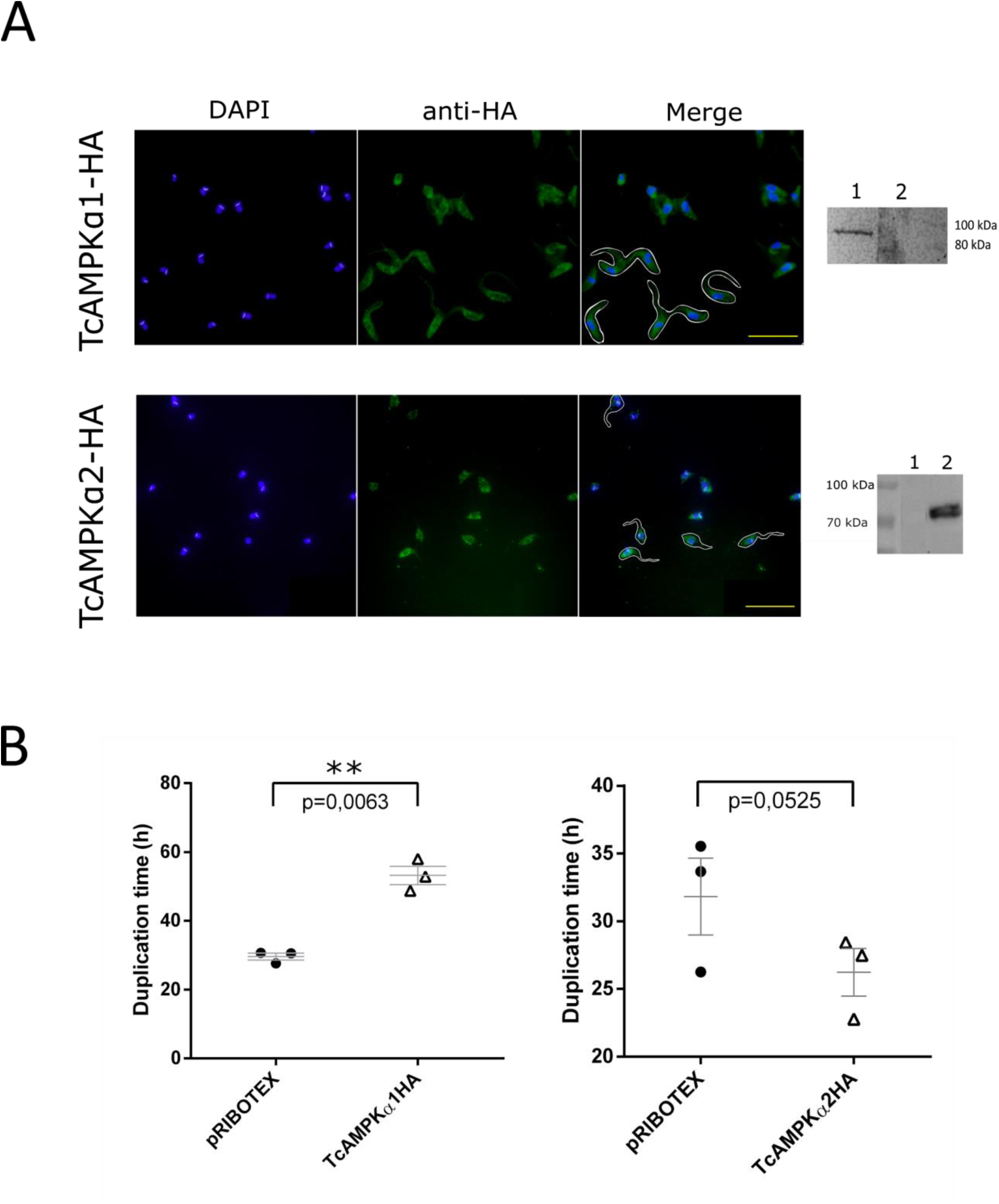
Over-expression of TcAMPKα isoforms in epimastigotes leads to antagonistic effects in proliferation. (A) Immunofluorescence images showing the cytoplasmic expression of the constructs. In the Merge quadrants some cells were contoured to illustrate the morphology of the parasites. Scale bar = 50 μm. To the right, western blots confirm tagged protein expression. (B) Duplication times of epimastigote cultures during the exponential phase of the growth curve (between day 3 and day 7) (n = 3). Error bars represent the Standard Error of the Mean and comparison was analyzed by a t-test. ** p value < 0.01

Cellular localization was similar for both proteins, distributed in the cytosol in a granulated pattern, which could point to a partial association with small organelles such as glycosomes or acidocalcisomes (Figure 5A). Parasites transfected with the empty vector or non-tagged construct didn’t show any fluorescent signal when revealed with anti-HA antibody.

We assessed the growth curves of the overexpressing cultures in regular conditions. Epimastigotes were counted in a Neubauer chamber for up to 10 days after initial dilution. Quantifications made between day 3 and 7 were considered as the exponential phase to evaluate the duplication time. Results show that overexpression of each isoform of the catalytic subunit causes antagonistic effects on proliferation (Figure 5B). Overexpression of TcAMPKα1-HA presents deleterious effects on epimastigotes, increasing duplication time in a statistically significant manner. These cultures didn’t reach maximum density and eventually arrested proliferation and died. Overexpression of TcAMPKα2-HA, instead, led to a slight decrease in duplication time during the exponential phase of the growth curve. These epimastigotes didn’t show any other effect on phenotype and reached normally the maximum density.

These results suggest that the TcAMPK complexes containing each catalytic subunit isoform could play different roles in the progression of the parasite life cycle.

### Relationship between TcAMPK and autophagic response in epimastigote cells

Since AMPK is involved in autophagy in several eukaryotic organisms in response to different kind of stresses, and considering the essential role of autophagy in the progression of *T. cruzi* life cycle, we sought to investigate if TcAMPK is capable of modulating autophagosome formation. To this end, we used Monodansylcadaverine (MDC), an acidotropic fluorescent dye that binds specifically to autophagosome membranes. After starving epimastigotes for 17 h in PBS, we measured MDC incorporation. We evaluated this parameter in wild type epimastigotes and the TcAMPKα2-HA overexpressing line.

Our results show that the overexpression of TcAMPKα2 leads to a higher autophagic capacity (Figure 6). The overexpressing epimastigotes can reach higher levels of MDC incorporation. Although marked inter-assay variations were observed, higher levels of MDC incorporation could always be registered in overexpressing parasites compared to WT ones. The intrinsic variations of the assay could be explained by the cyclic nature of autophagy itself, which will be further discussed in the Discussion section.

**Figure 6.**
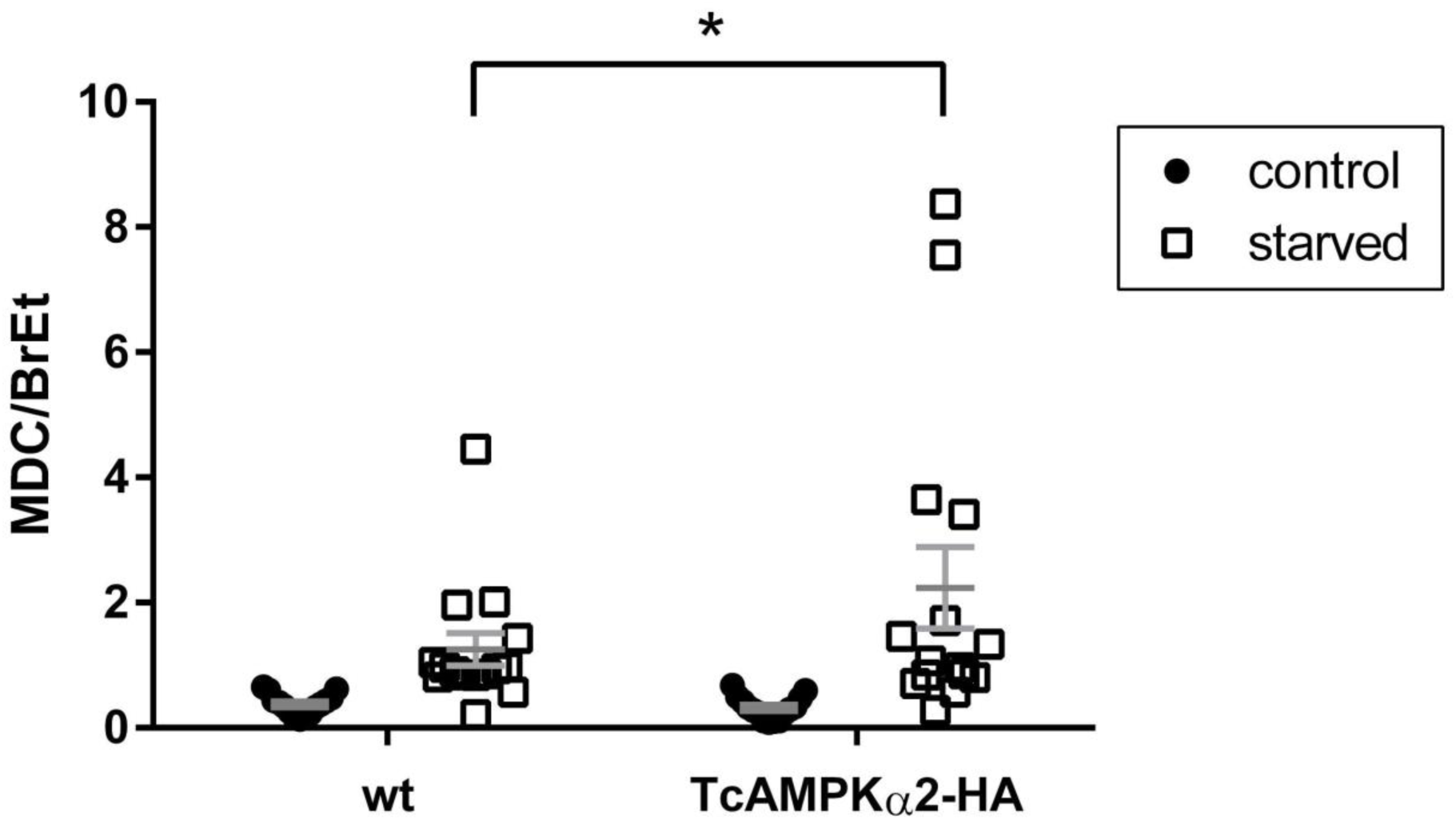
TcAMPKα2 overexpressing epimastigotes have an increased autophagic capacity. MDC incorporation was measured for wild type and overexpressing epimastigotes after 17 h incubation in either LIT medium (control) or PBS (starved). MDC values were normalized to Ethidium Bromide (BrEt) incorporation. MDC/BrEt ratio is expressed in arbitrary units. A t-test analysis of the starvation condition between the two strains shows a significant difference (n = 15). There were no significant differences in the control conditions. Error bars represent the Standard Error of the Mean.

## Discussion

*Trypanosoma cruzi* needs to precisely control energy homeostasis, nutrient availability, proliferation and differentiation in order to respond to changes in the environment and simultaneously preserve the viability of the host. In this sense, this parasite maintains a crucial coordination between these processes to achieve a host-parasite interaction and the permanence of the disease. Here, we report for the first time, the presence of two different AMPK complexes in *T. cruzi* and describe novel functions for TcAMPK as a key factor in the regulation of autophagy and proliferation, leading to a link between nutrient sensing, cell cycle and stage differentiation.

Our results show that *T. cruzi* expresses different AMPK complexes, fully capable of kinase activity on a specific AMPK substrate and actively regulated in the epimastigote stage, as shown by Thr172 phosphorylation and protein kinase activity under different treatments.

Yeast complementation exposed that every *T. cruzi* subunit has the necessary protein domains to function as part of an AMPK complex. Previous studies have shown that alpha subunits that cannot interact with the AMPK regulatory subunits, could be able to keep a basal protein kinase activity (5). Therefore, for these *T. cruzi* subunits, we could only establish a Snf1 function in yeast that restores their ability to use alternative carbon sources, although we cannot confirm glucose repression or interaction with the regulatory subunits. It is yet to be confirmed that all *T. cruzi* AMPK subunits interact with each other forming a heterotrimeric complex. Regarding this last concern, Saldivia et. al. had already demonstrated by co-immunoprecipitations, that both TbAMPK complexes, containing α1 or α2 subunits, are present in the bloodstream stage (12).

It is well known that AMPK in other eukaryotic organisms is regulated both allosterically and by post-translational modifications. The best defined mechanisms of AMPK activation are phosphorylation at Thr172 of the α-subunit and by AMP and/or adenosine diphosphate (ADP) binding to γ-subunit (28). Here, we demonstrated that TcAMPK could be regulated by phosphorylation at the Thr of its activation loop. Western blots using the Phospho-AMPKα(Thr172) antibody show two bands in epimastigotes extracts which sizes correspond with those of their theoretical molecular weights and are also similar to the weight of the *T. brucei* catalytic subunits. Another outstanding feature of the TcAMPKα2 subunit is the presence of a phosphorylatable Serine, instead of a Threonine, in the activation loop. To our knowledge, this is the only organism presenting this characteristic. The phosphorylation pattern of TcAMPKα subunits can be modified by nutritional stress and treatment with AMPK activators. To our knowledge, this is the only organism presenting this characteristic, and we hypothesize that it could differentially influence the upstream regulation of the complexes containing this subunit. These results point to the existence of at least one heterotrimeric complex in *T. cruzi*, which is able to sense AMP analogs and be activated by phosphorylation at the activation loop (in both alpha subunits) in consequence.

In order to assess the relative influence of AMPK activation on parasite metabolism, we used epimastigotes of *T. cruzi* overexpressing each of the catalytic subunits and found that this resulted in phenotypes associated with proliferation and nutritional stress response. Interestingly, these over-expressions had antagonistic effects on proliferation progression. TcAMPKα1 over-expression hindered proliferation, resulting in longer duplication times and in the inability to reach maximum parasite density, leading to eventual loss of the cultures. This inability to maintain stable transgenic cultures for TcAMPKα1 may be due to the fact that overexpressing the enzyme is detrimental for the progression of the cell cycle, possibly because parasites cannot cope with the induced metabolic stress. On the other side, the overexpression could be affecting cell cycle progression, leading to an arrest effect in proliferation. This in turn could be promoting the differentiation process (metacyclogenesis) that occurs in the final compartment of the intestinal tract (the posterior region of the small intestine and the rectum) (29). Over the years, several factors have been implicated to influence metacyclogenesis, such as osmolarity (30), the initial pH of the media (31), the carbon source availability (32) and more recently, autophagy (26,33,34). However, the molecular bases of the morphogenetic alterations necessary and sufficient to elicit parasite differentiation remain to be fully elucidated. It would be interesting, through other techniques such as the use of inducible vectors, to assess through which mechanism the constitutive overexpression of TcAMPKα1 causes the parasites to divide more slowly resulting in non-viable cell cultures. On the other hand, TcAMPKα2 over-expression led to shorter duplication times during the exponential growth phase. In other species, AMPK complexes have been shown to have antagonistic effects even with fewer amino acid sequence differences. This was also observed in *H. sapiens*, where AMPKα1 subunit has been classified as oncogene, because of its higher expression in some types of cancer cells, while AMPKα2 seems to function as a tumor suppressor instead (35). The authors state that these differences in function cannot be fully explained by differences in substrate specificity, but are rather determined by residues modified by regulatory upstream proteins, affecting AMPK activity and localization. In *T. cruzi*, sequence differences are more extensive between alpha subunits, and it would be important to evaluate if this affects substrate specificity and how, in addition to its regulation. Furthermore, epimastigotes over-expressing TcAMPKα2 were able to reach higher autophagic activity during prolonged nutritional stress, in comparison with control lines. Autophagy is an essential process in the progression of the life cycle of trypanosomatids (26,36,37) and AMPK is known to be involved in these processes in an ancestral and conserved manner (38–43). In this case, TcAMPKα2 would induce the autophagy process to cope with the nutritional stress caused by the absence of glucose as an energy source. Moreover, this coincides with an increase in AMPK activity in epimastigotes in the absence of glucose. The fact that TcAMPKα2 overexpression leads to an acceleration in cell duplication during the exponential phase can be interpreted as a role of the TcAMPKα2 subunit in the rapid consumption of glucose and maintenance of proliferation while there is availability of this carbon source, similarly to what would happen right after the insect vector takes a blood meal. After the insect hasn’t fed in several weeks, amino acids are the most abundant carbon source in the midgut and parasite density increases significantly. This leads to an arrest in proliferation, and TcAMPKa1 could then play a role in the initiation of metacyclogenesis. Its suggested role in proliferation and differentiation shows a possible neofunctionalization, involving metabolic pathways unique to these organisms. These results point to the presence of two AMPK complexes with distinct functions, although their exact roles in the parasite life cycle are yet to be determined.

Additionally to the role of AMPK in the rapid response and adaptation to stress, through the phosphorylation of downstream effectors, it also plays a part in the modulation of gene expression by phosphorylating transcription factors (44). In kinetoplastids, gene transcription is poorly modulated, there are no promoter sequences and transcription is carried out as polycistronic units processed by trans-splicing (45). mRNA transcription is then regulated by modifying these molecules stability, localization and translationality. For that reason, RNA Binding Proteins (RBPs) and RNA regulons have shown to have essential roles in different cellular processes in these organisms, such as differentiation and infectivity (46,47), and responses to nutrient and oxygen availability (48,49). One of our hypotheses regarding new roles for the AMPK complexes in trypanosomatids relates to a conserved capacity of this complex to induce gene expression by interacting with the RBPs and influencing regulon activation. Preliminary bioinformatic searches have shown some RBPs in *T. cruzi* and *T. brucei* that present the conserved sequences for AMPK phosphorylation, which makes them potential AMPK substrates. Furthermore, Saldivia et. al. identified the *T. brucei* RBP27 as an interactor of one of the TbAMPK complexes when co-immunoprecipitated. AMPK could function as the ancestral link between environmental conditions sensing and rapid and long term cellular response through these RBPs and other novel substrates.

To the extent of our knowledge this work brings the first evidence on a role for the AMPK complex in trypanosomatids during nutritional stress response. Our results suggest that the novel AMPK complex present in trypanosomatids was generated by AMPKα gene duplication in a common ancestor, preceding speciation, even though more experimental data must be generated to confirm this observation.

The finding in *T. cruzi* of a novel AMPK complex, which evolved incorporating unique characteristics, points out this pathway as a possible chemotherapy target for Chagas’ disease. Combining the background knowledge on AMPK complexes and the available compound libraries to affect its enzymatic activity, our novel TcAMPK complex finding makes it a promising target for therapeutic drug repositioning.

## Acknowledgments

We thank Dr. Martin G. Schmidt for generously sharing the yeast conditional mutants. We are grateful to Dr. Josefina Ocampo and Dr. Salomé Vilchez Larrea (INGEBI, Buenos Aires, Argentina) for helpful comments on the manuscript. This work was supported by the Consejo Nacional de Investigaciones Científicas y Técnicas (PIP 2013-00351); Departamento de Fisiología, Biología Molecular y Celular, Facultad de Ciencias Exactas y Naturales, Universidad de Buenos Aires (UBACyT 2014-2017, Nro 00155BA); and Agencia Nacional de Promoción Científica y Tecnológica (PICT 2013–2015, PICT 2015-0898 and PICT 2017-2125). G.D.A. and A.C.S. are members of the Research Career of Consejo Nacional de Investigaciones Científicas y Técnicas, T.S. is a fellow from the same institution. The funders had no role in study design, data collection and analysis, decision to publish, or preparation of the manuscript.

