## Supplemental Figure 1 for "AMP-activated protein kinase: A key enzyme to manage nutritional stress responses in parasites with complex life cycles"

### Supplementary Figure 1

Mass spectrometry analysis of epimastigotes protein extract to detect TcAMPKα subunits endogenous expression and its phosphorylation. (A) The central column of the table depicts the sequence of the peptide constructed from the protein fragments detected. This peptide corresponds to the activation loop of the TcAMPKα2, with a Serine in the position of the more conventional Thr172. Red and Blue numbers on the table are molecular weights of the peptides detected in the mass spectrometer, while black numbers are obtained by digital simulation of the protein digestion by the same proteases. (B) Spectra of the peptide sequence reconstructed in A. In the green circle is the peptide providing proof of the Serine phosphorylation.

A

| #1 | $b^{+}$ | $b^{2+}$ | $b^{3+}$ | Seq. | $y^{+}$ | $y^{2+}$ | $y^{3+}$ | #2 |
|---|---|---|---|---|---|---|---|---|
| 1 | 116.03422 | 58.52075 | 39.34959 | D | | | | 23 |
| 2 | 230.07715 | 115.54221 | 77.36390 | N | 2392.05267 | 1196.52997 | 798.02241 | 22 |
| 3 | 359.11974 | 180.06351 | 120.37810 | E | 2278.00974 | 1139.50851 | 760.00810 | 21 |
| 4 | 506.18815 | 253.59772 | 169.40090 | F | 2148.96715 | 1074.98721 | 716.99390 | 20 |
| 5 | 619.27222 | 310.13975 | 207.09559 | L | 2001.89873 | 1001.45301 | 667.97110 | 19 |
| 6 | 690.30933 | 345.65830 | 230.77463 | A | 1888.81467 | 944.91097 | 630.27641 | 18 |
| 7 | 857.30769 | 429.15748 | 286.44075 | S-Phospho | 1817.77756 | 909.39242 | 606.59737 | 17 |
| 8 | 944.33972 | 472.67350 | 315.45142 | S | 1650.77920 | 825.89324 | 550.93125 | 16 |
| 9 | 1104.37037 | 552.68882 | 368.79497 | Carbamidometl | 1563.74717 | 782.37722 | 521.92057 | 15 |
| 10 | 1161.39183 | 581.19955 | 387.80213 | G | 1403.71652 | 702.36190 | 468.57702 | 14 |
| 11 | 1248.42386 | 624.71557 | 416.81280 | S | 1346.69506 | 673.85117 | 449.56987 | 13 |
| 12 | 1345.47662 | 673.24195 | 449.16373 | P | 1259.66303 | 630.33515 | 420.55919 | 12 |
| 13 | 1459.51955 | 730.26341 | 487.17803 | N | 1162.61026 | 581.80877 | 388.20827 | 11 |
| 14 | 1622.58288 | 811.79508 | 541.53248 | Y | 1048.56734 | 524.78731 | 350.19396 | 10 |
| 15 | 1693.61999 | 847.31363 | 565.21152 | A | 885.50401 | 443.25564 | 295.83952 | 9 |
| 16 | 1764.65711 | 882.83219 | 588.89055 | A | 814.46689 | 407.73709 | 272.16048 | 8 |
| 17 | 1861.70987 | 931.35857 | 621.24147 | P | 743.42978 | 372.21853 | 248.48144 | 7 |
| 18 | 1990.75246 | 995.87987 | 664.25567 | E | 646.37702 | 323.69215 | 216.13052 | 6 |
| 19 | 2103.83653 | 1052.42190 | 701.95036 | I | 517.33442 | 259.17085 | 173.11633 | 5 |
| 20 | 2216.92059 | 1108.96393 | 739.64505 | L | 404.25036 | 202.62882 | 135.42164 | 4 |
| 21 | 2303.95262 | 1152.47995 | 768.65572 | S | 291.16630 | 146.08679 | 97.72695 | 3 |
| 22 | 2360.97408 | 1180.99068 | 787.66288 | G | 204.13427 | 102.57077 | 68.71627 | 2 |
| 23 | | | | K | 147.11280 | 74.06004 | 49.70912 | 1 |

B

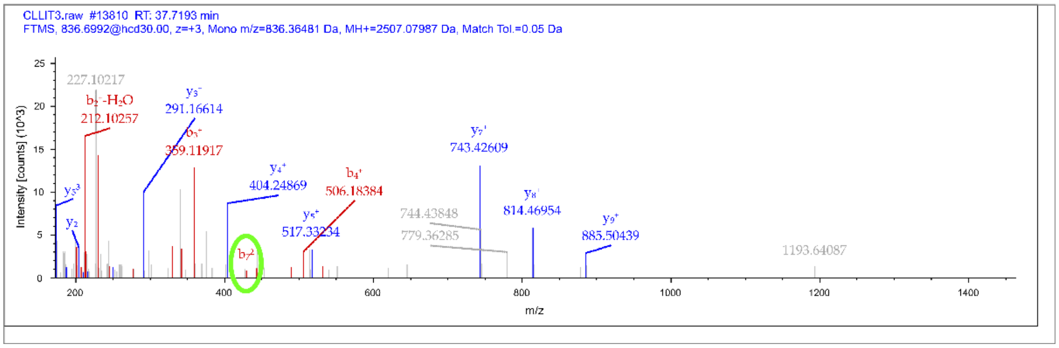
